# GABAergic synaptic scaling is triggered by changes in spiking activity rather than AMPA receptor activation

**DOI:** 10.1101/2023.03.08.531789

**Authors:** Carlos Gonzalez-Islas, Zahraa Sabra, Ming-fai Fong, Pernille Bülow, Nicholas Au Yong, Kathrin Engisch, Peter Wenner

## Abstract

Homeostatic plasticity represents a set of mechanisms that are thought to recover some aspect of neural function. One such mechanism called AMPAergic scaling was thought to be a likely candidate to homeostatically control spiking activity. However, recent findings have forced us to reconsider this idea as several studies suggest AMPAergic scaling is not directly triggered by changes in spiking. Moreover, studies examining homeostatic perturbations *in vivo* have suggested that GABAergic synapses may be more critical in terms of spiking homeostasis. Here we show results that GABAergic scaling can act to homeostatically control spiking levels. We found that perturbations which increased or decreased spiking in cortical cultures triggered multiplicative GABAergic upscaling and downscaling, respectively. In contrast, we found that changes in AMPAR or GABAR transmission only influence GABAergic scaling through their indirect effect on spiking. We propose that GABAergic scaling represents a stronger candidate for spike rate homeostat than AMPAergic scaling.

**Significance Statement:** The nervous system maintains excitability in order to perform network behaviors when called upon to do so. Networks are thought to maintain spiking levels through homeostatic synaptic scaling, where compensatory multiplicative changes in synaptic strength are observed following alterations in cellular spike rate. Although we demonstrated that AMPAergic synaptic scaling does not appear meet these criteria as a spike rate homeostat, we now show that GABAergic scaling could play this role. Here we present evidence that the characteristics of GABAergic scaling place it in an excellent position to be a spiking homeostat. This work highlights the importance of inhibitory circuitry in the homeostatic control of excitability. Further, it provides a point of focus into neurodevelopmental disorders where excitability is impaired.

## Introduction

Homeostatic plasticity represents a set of compensatory mechanisms that are thought to be engaged by the nervous system in response to cellular or network perturbations, particularly in developing systems ^1^. Synaptic scaling is one such mechanism where homeostatic compensations in the strength of the synapses onto a neuron occur following chronic perturbations in spiking activity or neurotransmitter receptor activation (neurotransmission) ^2^. Scaling is typically identified by comparing the distribution of miniature postsynaptic current (mPSC) amplitudes in control and activity-perturbed conditions. For instance, when spiking activity in cortical cultures was reduced for 2 days with the Na^+^ channel blocker TTX or the AMPA/kainate glutamate receptor antagonist CNQX, mEPSC amplitudes were increased ^2^. When first discovered, homeostatic synaptic scaling was thought to be triggered by the cell sensing its reduction in spike rate through reduced calcium entry into the cytoplasm. This was then believed to alter global calcium signaling cascades that led to increased AMPA receptor (AMPAR) insertion in a cell-wide manner such that all synapses increased synaptic strength multiplicatively based on each synapse’s initial strength ^3^. In this way excitatory synaptic strength was increased across all of the cell’s inputs in order to recover spiking activity without altering relative synaptic strengths resulting from Hebbian plasticity mechanisms. These criteria, sensing spike rate and adjusting synaptic strengths multiplicatively, thus established the expectations for homeostatic synaptic scaling and were consistent with the idea that AMPAergic scaling could be a spike rate homeostat.

More recent work has demonstrated that AMPAergic synaptic scaling is more complicated than originally thought. First, studies have now shown that increases in mEPSC amplitudes or synaptic glutamate receptors often do not follow a simple multiplicative function ^4, 5^. Rather, these studies show that changes in synaptic strength at different synapses exhibit different scaling factors, arguing against a single multiplicative scaling factor that alters synaptic strength globally across the cell. Second, AMPAergic scaling triggered by receptor blockade can induce a synapse-specific plasticity rather than a cell-wide plasticity. Compensatory changes in synaptic strength were observed in several studies where neurotransmission at individual synapses was reduced ^6–9^. This synapse-specific plasticity would appear to be cell-wide if neurotransmission at all synapses were reduced as occurs in the typical pharmacological blockades that are used to trigger scaling. Regardless, this would still be a synapse specific plasticity, determined at the synapse, rather than the cell sensing its lowered spiking activity through global calcium levels. Finally, several different studies now suggest that reducing spiking levels in neurons is not sufficient to trigger AMPAergic upscaling and therefore bring into question its role as a spike rate homeostat. Forced expression of a hyperpolarizing conductance reduced spiking of individual cells but did not trigger AMPAergic scaling ^10^. Further, optogenetic restoration of culture-wide spiking in the presence of AMPAergic transmission blockade triggered AMPAergic scaling that was indistinguishable from that of cultures where AMPAR block reduced spiking (no optogenetic restoration of spiking) ^11^. Most studies that separate the importance of cellular spiking from synapse-specific transmission suggest that AMPAergic scaling is triggered by changes in neurotransmission, rather than a cell’s spiking activity ^9–12^. While transmission-dependent AMPAergic scaling appears to be more commonly observed, there are two studies that suggest that alterations in AMPAergic synaptic strength can occur following alterations in spiking in individual cells – AMPA receptor accumulation following blockade of spiking at the soma in cortical cultures ^13^ and reduced mEPSC amplitude following optogenetic activation of individual cells in hippocampal cultures ^14^.

Because the pharmacological perturbations that trigger AMPAergic upscaling also result in GABAergic downscaling, it is assumed that they have common triggers. Therefore, in the current study we tested this possibility. Homeostatic regulation of GABAergic miniature postsynaptic current (mIPSC) amplitude was first shown in excitatory neurons following network activity perturbations ^15^. Similar to AMPAergic upscaling, chronic perturbations in AMPAR or spiking activity triggered mIPSC downscaling through compensatory changes in the number of synaptic GABA_A_ receptors ^15–19^. However, the sensing machinery for triggering GABAergic scaling may be distinct from that of AMPAergic scaling ^20^. Further, GABAergic plasticity does appear to be a key player in the homeostatic response *in vivo*, as many different studies have shown strong GABAergic compensations following somatosensory, visual, and auditory deprivations ^21-25^. In addition, these homeostatic GABAergic responses precede and can outlast compensatory changes in the glutamatergic system. Here we describe that GABAergic scaling is triggered by changes in spiking levels rather than changes in AMPAergic or GABAergic neurotransmission, that GABAergic scaling is expressed in a multiplicative manner, and could contribute to the homeostatic recovery of spiking activity. Our results suggest that GABAergic scaling could serve as a homeostat for spiking activity.

## Results

### TTX and AMPAR blockade triggered a non-uniform scaling of AMPA mPSCs

Previously we have shown that blocking spike activity in neuronal cultures triggered AMPAergic scaling in a non-uniform or divergent manner, such that different synapses scaled with different scaling ratios ^4, 26^. Importantly, these results were consistent across independent studies performed in three different labs using rat or mouse cortical cultures, or mouse hippocampal cultures. We quantitatively evaluated scaling in the following manner. We rank-ordered mEPSC amplitudes (smallest to largest) for both control and TTX-treated cultures and then divided the TTX rank ordered amplitude by the corresponding control rank ordered amplitude (e.g. smallest TTX amplitude divided by smallest control amplitude, etc.) and plotted these ratios for all such comparisons ^4, 26^. Previously, scaling had been thought to be multiplicative, meaning all mPSC amplitudes were altered by a single multiplicative factor. If true for AMPAergic scaling, then our ratio plots should have produced a horizontal line at the scaling ratio. However, we found that ratios progressively increased across at least 75% of the distribution of amplitude ratios. Still, it was unclear whether this was true for all forms of AMPAergic scaling triggered by different forms of activity blockade. Therefore, we repeated this analysis on the data from our previous study ^11^, but now on AMPAergic scaling produced by blocking AMPAR neurotransmission (40µM CNQX), rather than TTX. We found that the scaling was non-uniform and replicated the scaling triggered by TTX application (Supplemental Figure 1). There was an abrupt increase in the ratio from 1 to ∼1.2 (steeper slope) over the first 1-2% of the data, consistent with an error caused by the detection threshold (as shown in simulations of a threshold issue in ^4^). However, ratios then increased over the vast majority of the data from 1.2 to 1.5 more slowly, and this represented the magnitude of homeostatic plasticity with increasing mEPSC amplitude. The results suggest that AMPAergic scaling produced by blocking glutamatergic transmission or spiking in culture was not multiplicative, but rather different synapses increased by different scaling factors. Further, the similarity of scaling ratio plots following either action potential or AMPAergic blockade is consistent with the idea that they are mediated by similar mechanisms.

### TTX and AMPAR blockade reduced both spiking and GABAergic mIPSC amplitude

Previously we made the surprising discovery that AMPAergic upscaling in rat cortical cultures was triggered by a reduction in AMPAR activation rather than a reduction in spiking activity ^11^. Here we tested whether GABAergic scaling was dependent on AMPAR activation or a different trigger, by changes in spiking activity levels. We plated E18 mouse cortical neurons on 64 channel planar multi-electrode arrays (MEAs) and allowed the networks to develop for ∼14 days *in vitro* (DIV), a time point where most cultures develop a network bursting behavior (Figure 1A, Supplemental Figure 2) ^27^. We used a custom written Matlab program that was able to detect and compute overall spike rate and burst frequency (Supplemental Figure 2, see methods). We again found that TTX abolished bursts and spiking activity (n=2, Supplemental Figure 3). On the other hand, AMPAR blockade (20µM) merely reduced bursts and spiking, with a greater effect on bursting. Similar to our findings in rat cortical cultures ^11^, CNQX dramatically reduced burst frequency and maintained this reduction for the entire 24hrs of treatment (Figure 1B). Overall spike frequency was also reduced in the first 6 hours, but then recovered over the 24 hour drug treatment (Figure 1C). While overall spiking was recovered, we did note that this was highly variable, with some cultures recovering minimally. Following AMPAergic blockade bursts continued in these cultures, likely due to NMDAergic neurotransmission as shown previously ^11^.

**Figure 1.**
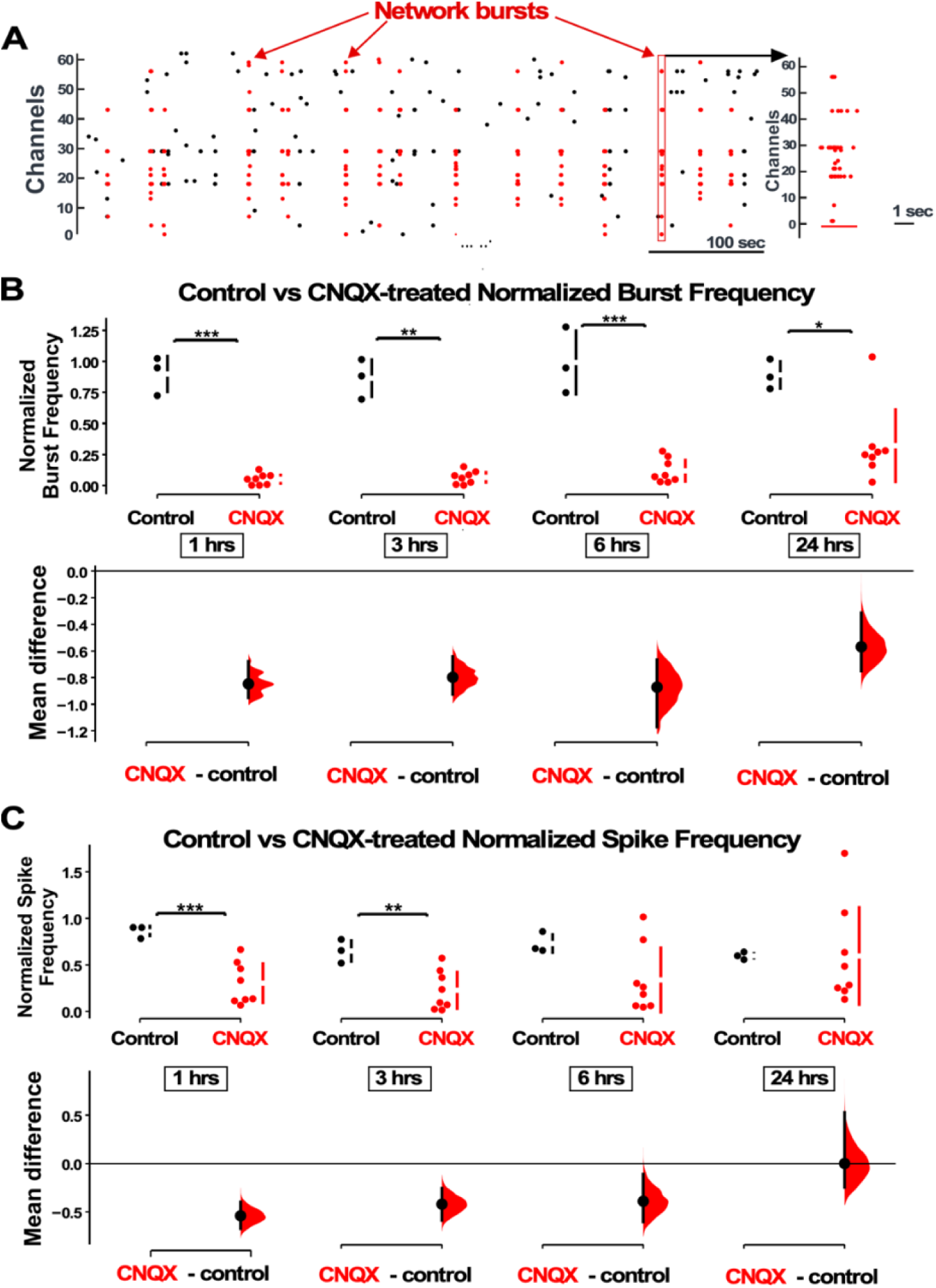
AMPAergic blockade reduces burst frequency and overall spike rate. **A**) Network bursts can be identified by detected spikes (red dots) time-locked in multiple channels of the MEA (Y axis). One burst (red rectangle) is expanded in time and shown in the raster plot on the right. **B**) The normalized burst rate is shown in control cultures and following application of CNQX for 24 hrs. **C**) Average overall spike frequency is compared for CNQX-treated cultures and control unstimulated cultures at 1hr, 3hrs, 6hrs (p=0.104), and 24hrs (p=0.982) after addition of CNQX or vehicle. The mean differences at different time points are compared to control and displayed in Cumming estimation plots. Significant differences denoted by * p ≤ 0.05, ** p ≤ 0.01, *** p ≤ 0.001. Recordings from single cultures (filled circles, control n = 3 cultures, CNQX n = 8 cultures), where mean values (represented by the gap in the vertical bar) and SD (vertical bars) are plotted on the upper panels. Mean differences between control and treated groups are plotted on the bottom panel, as a bootstrap sampling distribution (mean difference is represented by a filled circles and the 95% CIs are depicted by vertical error bars).

In order to examine the possibility that compensatory changes in GABAergic synaptic strength could have contributed to the recovery of the network spiking activity we assessed synaptic scaling by measuring mIPSC amplitudes in pyramidal-like neurons in a separate set of cortical cultures plated on coverslips. We found that both activity blockade with TTX, and AMPAergic blockade with CNQX, triggered a dramatic compensatory reduction in mIPSC amplitude compared to control (untreated) cultures (Figure 2A). Even though TTX completely abolished spiking, while CNQX only reduced spiking, both treatments triggered a similar reduction in average mIPSC amplitude. In order to more carefully compare the GABAergic scaling that is triggered by TTX and CNQX mechanistically, we created scaling ratio plots as described above ^4^. In Figure 2B we show that TTX-induced and CNQX-induced scaling does produce a largely multiplicative downscaling with a scaling factor around 0.5. This is consistent with the idea that the mechanisms of GABAergic scaling were similar following activity or AMPAergic blockade. We noticed that the first mIPSC ratios started near one and within 50 ratios came down to the 0.5 level (Figure 2B). This is likely due to the smallest drug-treated mIPSCs falling below our detection cutoff of 5 pAs ^4^. On the other hand, the largest mIPSCs trended above 0.5, consistent with the possibility that a small portion of the mIPSCs may not scale uniformly. Together, these results are consistent with the idea that either spiking or reduced AMPA receptor activation could trigger the GABAergic downscaling since TTX or CNQX would reduce both.

**Figure 2.**
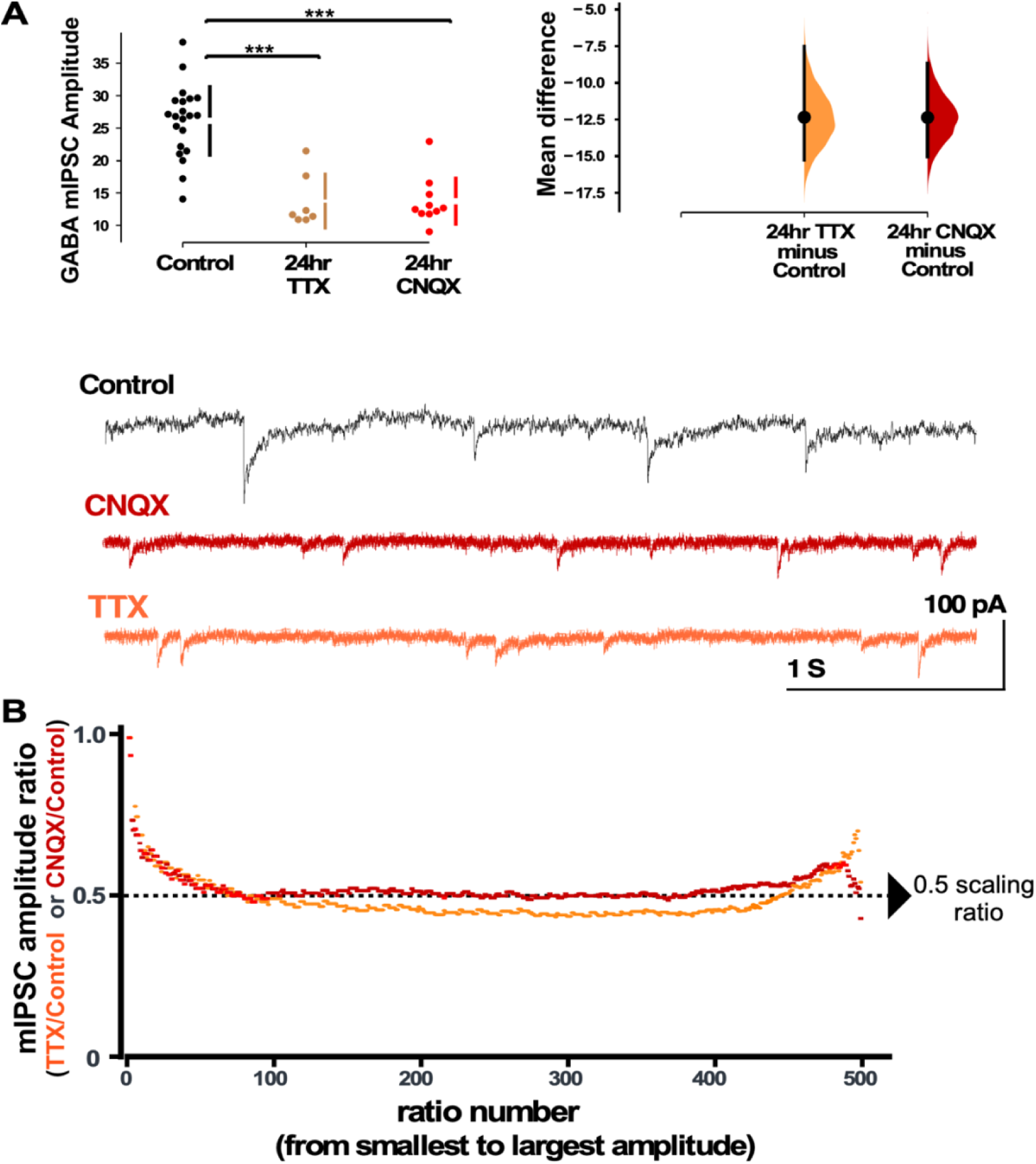
Both activity and AMPAR blockade cause a reduction in mIPSC amplitudes that appear to scale down. **A**) CNQX and TTX produce a reduction in average amplitude of mIPSCs as shown in the scatter plots (control - n=21 from 10 cultures, TTX - n=7 from 3 cultures, CNQX - n=10 from 6 cultures). The mean differences are compared to control and displayed in Cumming estimation plots. Significant differences denoted by *** p ≤ 0.001. GABAergic mPSC amplitudes from single neurons (filled circles), where mean values (represented by the gap in the vertical bar) and SD (vertical bars) are plotted on the panels to the left. Mean differences between control and treated groups are plotted on the panel to the right, as a bootstrap sampling distribution (mean difference is represented by a filled circles and the 95% CIs are depicted by vertical error bars). Example traces showing mIPSCs are shown below. **B**) Scaling ratio plots show the relationship of mIPSC amplitudes from treated cultures compared to untreated cultures. All recordings taken from cultured neurons plated on coverslips, not MEAs.

### Optogenetic restoration of spiking in the presence of AMPAR blockade prevented GABAergic downscaling

In order to separate the importance of spiking levels from AMPAR activation in triggering GABAergic downscaling we blocked AMPARs while restoring spike frequency. Cultures were plated on the MEA and infected with ChR2 under the human synapsin promoter on DIV 1. Experiments were carried out on ∼ DIV14, when cultures typically express network bursting. Baseline levels of spike frequency were measured in a 3-hour period before the addition of 20 µM CNQX (Figure 3A). We then used a custom written TDT Synpase software that triggered a brief (50-100ms) activation of a blue light photodiode to initiate bursts (see methods, Figure 3B) whenever the running average of the firing rate fell below the baseline level, established before the addition of the drug. In this way we could optically initiate bursts that largely occurred after the blue light was off. These optically-induced bursts look very similar to the spontaneously occurring pre-drug bursts and this largely restored the spike rate to pre-drug values (Figure 3B). We used 20 µM CNQX to block AMPARs, instead of the 40 µM concentration that we used in the previous study ^11^ because 40 µM CNQX severely impaired our ability to optogenetically restore spiking activity in these cultures.

**Figure 3.**
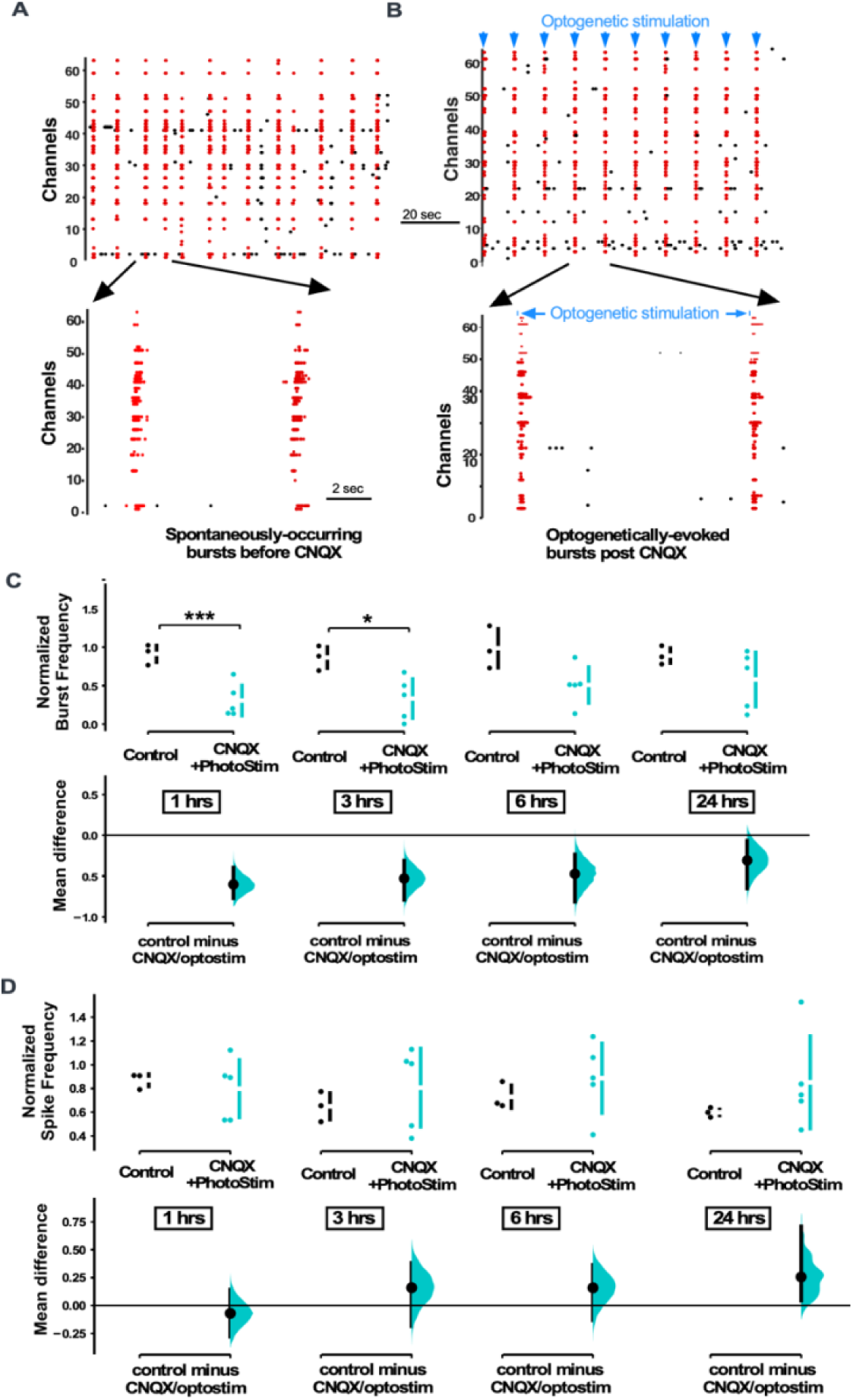
MEA recordings show that optogenetic stimulation restores spiking activity in cultures treated with CNQX. **A**) Spontaneously-occurring bursts of spiking are identified (synchronous spikes/red dots). Expanded version of raster plot highlighting 2 bursts is shown below. **B**) Same as in A, but after CNQX was added to the bath and bursts were now triggered by optogenetic stimulation (blue line shows duration of optogenetic stimulation). **C**) Average burst rate is compared for CNQX-treated cultures with optogenetic stimulation (n=5 cultures) and control unstimulated cultures (n=3 cultures) at 1hr, 3hrs, 6hrs (p=0.056), and 24hrs (p=0.379) after addition of CNQX or vehicle (same control data presented in Figure 1). **D**) Average overall spike frequency is compared for CNQX-treated cultures with optogenetic stimulation and control unstimulated cultures at 1hr (p=0.612), 3hrs (p=0.489), 6hrs (p=0.449), and 24hrs (p=0.22) after addition of CNQX or vehicle. Control data is same as presented in Figure 1. The mean differences at different time points are compared to control and displayed in Cumming estimation plots. Significant differences denoted by * p ≤ 0.05, *** p ≤ 0.001. Recordings from single cultures (filled circles), where mean values (represented by the gap in the vertical bar) and SD (vertical bars) are plotted on the upper panels. Mean differences between control and treated groups are plotted on the bottom panel, as a bootstrap sampling distribution (mean difference is represented by a filled circles and the 95% CIs are depicted by vertical error bars).

We have already established that bursts and spiking were reduced following the application of CNQX (Figure 1). However, when we optogenetically activated the cultures in the presence of CNQX we found that both the burst rate and spike frequency were increased compared to CNQX treatment alone, no optostimualtion (Supplemental Figure 4). Because the program was designed to maintain total spike frequency, photostimulation of CNQX-treated cultures did a relatively good job at recovering this parameter to control levels (Figure 3D). In fact, spike frequency was slightly, but not significantly, above control levels through the 24 hour recording period (Figure 3D). In our previous study we were able to establish that the optogenetically-evoked bursts in CNQX and even the pattern of individual unit spiking during the burst was restored to that of normally occurring bursts in the pre-drug condition ^11^. On the other hand, the program designed to control overall spike frequency through optostimulation in CNQX did not completely return burst frequency back to control levels (Figure 3C).

We next assessed mIPSC amplitudes using whole cell recordings taken from cultures plated on MEAs. After blocking AMPAR activation without optogenetic restoration of spiking activity, we found that mIPSC amplitudes were significantly reduced compared to control conditions (Figure 4A), as we had shown for CNQX treatment on cultures plated on coverslips (Figure 2A). Strikingly, when spiking activity was optogentically restored in the presence of CNQX for 24 hours we observed that mIPSCs were no different than control values (same as control, larger than CNQX only – Figure 4A). This result suggested that unlike AMPAergic upscaling, GABAergic downscaling was prevented if spiking activity levels were restored in the presence of AMPAR blockade. In order to compare scaling profiles we plotted the scaling ratios for these different treatments. Not surprisingly, we found that MEA-plated cultures treated with CNQX but given no optogenetic stimulation were similar to CNQX-treated cultures plated on coverslips (CNQX/control ∼ 0.5, Figure 4B vs Figure 2B). Ratio plots of cultures treated with CNQX where activity was restored optogenetically compared to controls demonstrated a fairly uniform relationship with a ratio of around 1 through most of the distribution suggesting the mIPSCs in these two conditions were similar and therefore unscaled (Figure 4B). Interestingly, the scaling ratio and the average mIPSC amplitudes in the optogenetically activated cultures were slightly larger than control mIPSCs which may be due to the slight increase in spiking in optogeneticallly stimulated cultures. Together, these results suggest that GABAergic downscaling was triggered by reductions in spiking activity, independent of AMPA receptor activation, and was multiplicative in that the vast majority of mEPSC amplitudes (∼95%) appeared to be reduced to ∼50%.

**Figure 4.**
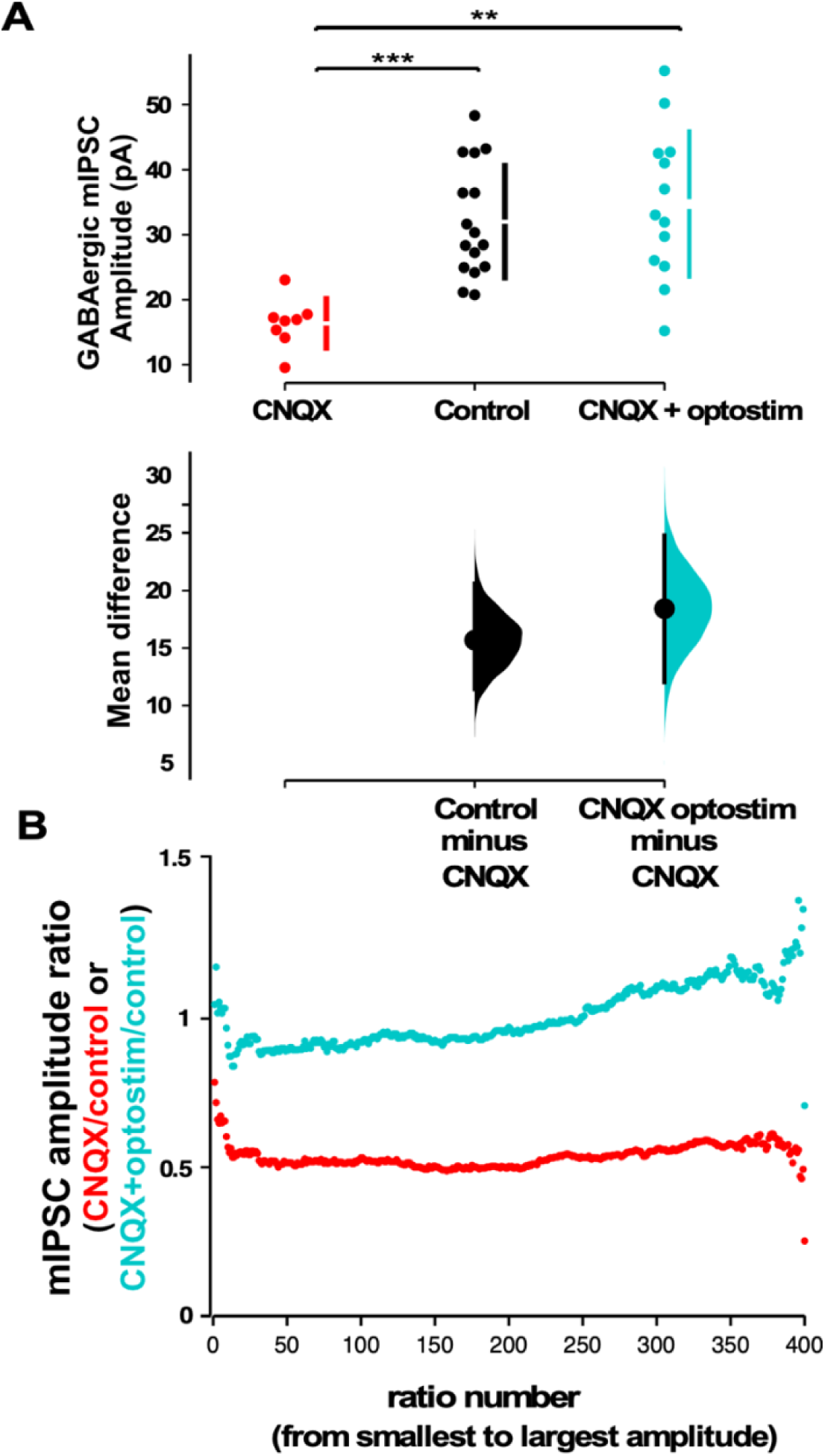
Optogenetic restoration of spiking activity in the presence of AMPAR blockade prevents GABAergic downscaling observed in CNQX alone. **A**) Scatter plots show AMPAR blockade triggers a reduction in mIPSC amplitude compared to controls that is prevented when combined with optogenetic stimulation (optostim, control - n=16 from 10 cultures, CNQX - n=8 from 4 cultures, CNQX/optostim - n=13 from 6 cultures). The mean differences are compared to control and displayed in Cumming estimation plots. Significant differences denoted by ** p ≤ 0.01, *** p ≤ 0.001. GABAergic mIPSC amplitudes from single neurons (filled circles), where mean values (represented by the gap in the vertical bar) and SD (vertical bars) are plotted on the upper panels. Mean differences between control and treated groups are plotted on the bottom panel, as a bootstrap sampling distribution (mean difference is represented by a filled circles and the 95% CIs are depicted by vertical error bars). **B**) Scaling ratio plots show largely multiplicative relationships to control values for both CNQX and CNQX + photostimulation treatments. Cultured neurons for these recordings were obtained from cells plated on MEAs (control, CNQX, and CNQX+optostim).

### Enhancement of AMPAR currents triggered GABAergic upscaling in a spike-dependent manner

While reductions in spiking activity triggered a GABAergic downscaling, it was less clear whether increases in spiking activity could trigger compensatory GABAergic upscaling. To test for such a possibility, we exposed the cultures to cyclothiazide (CTZ), an allosteric enhancer of AMPA receptors that also enhances spontaneous glutamate release ^11^. Due to the hydrophobic nature of CTZ it was necessary to dissolve it in ethanol, and used ethanol without CTZ as a control (final solution 1:1000 ethanol in Neurobasal). In addition to increasing AMPAR activation, CTZ application slightly increased overall spiking activity in our MEA-plated cultures in the first 3 hours of the drug, although this was quite variable (Figure 5A-B). The amplitude of mIPSCs in control cultures exposed to ethanol were no different than control cultures without ethanol (Figure 5C). We then treated coverslip-plated cultures with CTZ for 24 hours and found that this did indeed produce a compensatory increase in GABA mIPSC amplitude (Figure 5D). In our previous study we found that enhancing AMPAergic neurotransmission in the presence of activity blockade (CTZ + TTX) reduced AMPAergic upscaling compared to activity blockade alone (TTX) ^11^. Therefore, we tested whether enhancing AMPAergic neurotransmission in activity blockade (CTZ + TTX) altered GABAergic scaling induced by TTX alone (24 hrs). Here we found that the GABAergic downscaling following TTX was no different when AMPAergic neurotransmission was enhanced (CTZ + TTX, Figure 5D). To determine if these changes in mIPSC amplitude were of a multiplicative scaling nature we made ratio plots. This demonstrated that both CTZ increases and CTZ+TTX decreases in mIPSC amplitude were multiplicative and therefore represented scaling (Figure 5E, CTZ – scaling ratio of 1.5, CTZ+TTX - scaling ratio of 0.6). Further, the scaling ratio plot for CTZ + TTX looked similar to those of TTX alone (compare Figure 5E and 2B). These results showed a compensatory upward and downward GABAergic scaling, that more closely followed spiking activity compared to AMPAergic transmission.

**Figure 5.**
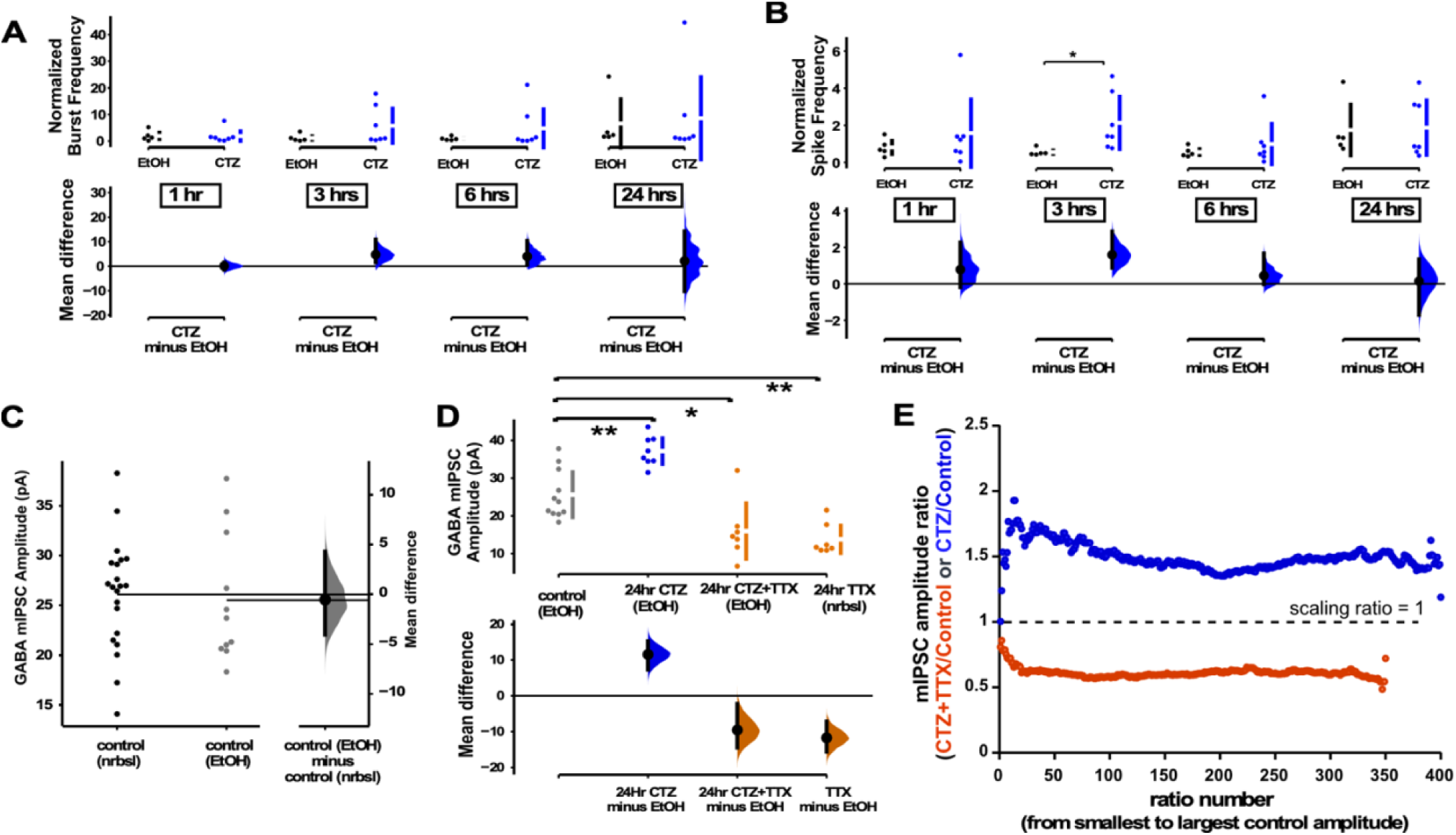
GABAergic upscaling was also triggered by CTZ and this was dependent on spiking activity. **A**) MEA recordings show that CTZ-treated cultures trended toward increases in normalized burst rate compared to control untreated cultures at 1hr (p=0.97), 3hrs (p=0.246), 6hrs (p=0.397), and 24hrs (p=0.894) after addition of CNQX (n=7) or vehicle (n=5). **B**) MEA recordings show that CTZ-treated cultures trended toward increases in normalized overall spike rate compared to control untreated cultures at 1hr (p=0.565), 3hrs, 6hrs (p=0.634), and 24hrs (p=0.92) after addition of CNQX or vehicle. **C**) Control cultures in Neurobasal (nrbsl) were compared with control cultures with ethanol (EtOH) dissolved in Neurobasal (1:1000). Amplitude of mIPSCs in different controls were no different (p=0.803, nrbsl - n=21 from 10 cultures, EtOH - n=11 from 3 cultures). **D**) CTZ treatment (dissolved in ethanol) led to an increase in mIPSC amplitude compared to ethanol control cultures (CTZ - n=8 from 3 cultures). CTZ combined with TTX (in ethanol) produced a reduction of mIPSC amplitude compared to controls (ethanol) that was no different than TTX (nrbsl) alone (CTZ+TTX - n=7 from 3 cultures, TTX - n=7 from 3 cultures is same data as shown in Figure 2A). The mean differences at different time points or conditions are compared to control and displayed in Cumming estimation plots. Significant differences denoted by * p ≤ 0.05, ** p ≤ 0.01. Recordings from single cultures (filled circles), where mean values (represented by the gap in the vertical bar) and SD (vertical bars) are plotted on the upper panels. Mean differences between control and treated groups are plotted on the bottom panel, as a bootstrap sampling distribution (mean difference is represented by a filled circles and the 95% CIs are depicted by vertical error bars). **E**) Scaling ratios show that both CTZ-induced increases and CTZ+TTX -induced decreases were multiplicative. All mIPSC amplitudes recorded from cultures plated on coverslips, not MEAs.

### Blocking GABAergic receptors for 24 hours triggered upscaling of GABAergic mIPSCs

The above results suggested that GABAergic scaling was more dependent on the levels of spiking activity. However, one alternative possibility was that these changes in GABA mPSCs were due to changes in GABAergic receptor activation, which could act as a proxy for activity levels. In this way, GABARs sense changes in spiking activity levels and directly trigger GABAergic scaling to recover activity. To address this possibility, we treated cultures with the GABA_A_ receptor antagonist bicuculline to chronically block GABAergic receptor activation while increasing spiking activity. If increased spiking activity is directly the trigger (not mediated through GABAR activity), then we would expect to see GABAergic upscaling. On the other hand, if GABAR activation is a proxy for spiking then blockade of these receptors would indicate low activity levels and we would expect a downscaling to recover the apparent loss of spiking. GABAR block produced an upward trend in both burst frequency (Figure 6A) and spike frequency (Figure 6B). We measured mIPSCs in a separate cohort of cultures plated on coverslips which were treated with bicuculline for 24 hours, and we observed GABAergic upscaling (Figure 6C). These results are consistent with previous work in hippocampal cultures that showed GABAergic upscaling following bicuculline treatment^18, 28^. We also assessed mIPSC frequency in all of the drug conditions but did not observe significant differences, possibly due to the tremendous variability of this feature (Supplemental Figure 5). Our results were consistent with the idea that direct changes in spiking activity, rather than AMPA or GABA receptor activation triggered compensatory GABAergic upscaling. The scaling ratio plots were again relatively flat, with a scaling ratio of around 1.5, suggesting a multiplicative GABAergic upscaling (Figure 6D) that was similar to CTZ-induced upward scaling (Figure 5E).

**Figure 6.**
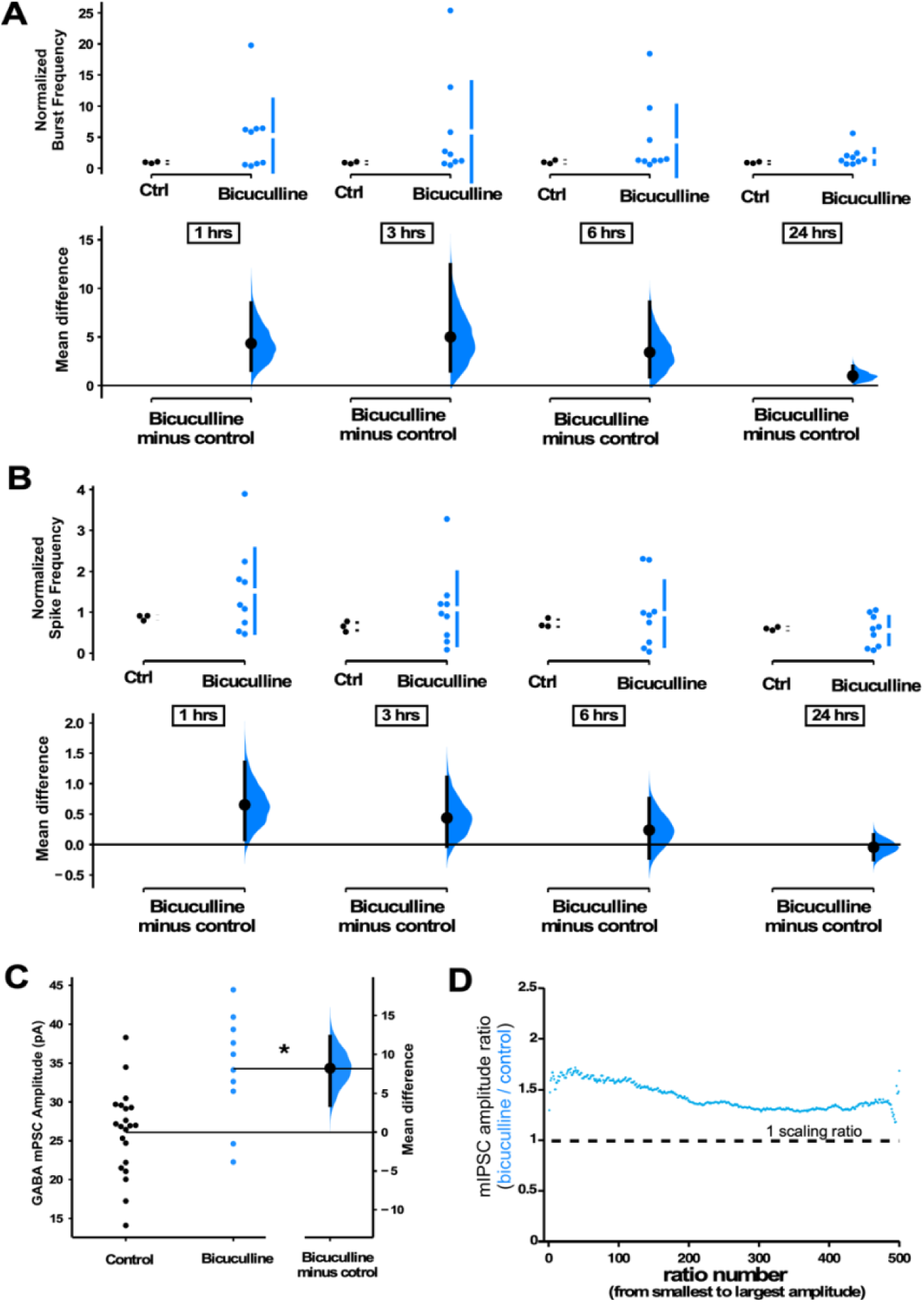
GABAergic upscaling is triggered by increased spiking activity rather than reduced GABAR activation. **A**) Bicuculline-treated cultures (24hrs) plated on MEA’s trended upward in normalized burst rate compared to control untreated cultures at 1hr (p=0.314), 3hrs (p=0.246), 6hrs (p=0.339), and 24hrs (p=0.339) after addition of bicuculline (n=9 cultures) or vehicle (n=3 cultures, same data as Figure 1). **B**) Bicuculline-treated cultures (24hrs) plated on MEA’s trended upward in normalized overall spike frequency compared to control untreated cultures at 1hr (p=0.358), 3hrs (p=0.462), 6hrs (p=0.734), and 24hrs (p=0.772) after addition of bicuculline or vehicle. Recordings from single cultures (filled circles), where mean values (represented by the gap in the vertical bar) and SD (vertical bars) are plotted on the upper panels. **C**) Bicuculline treatment (24hrs) produced an increase in mIPSC amplitudes (control n=21 form 10 cultures, bicuculline - n=10 from 4 cultures). The mean difference is compared to control and displayed in Cumming estimation plots. Significant difference denoted by * p ≤ 0.05. Recordings from single neurons (filled circles), and mean values (represented by the horizontal line). Control and treated group is plotted, as a bootstrap sampling distribution (mean difference is represented by a filled circles and the 95% CI is depicted by vertical error bar). **D**) Ratio plots for bicuculline-induced increase in mIPSCs exhibits a multiplicative profile. All mIPSC amplitudes recorded from cultures plated on coverslips, not MEAs.

### The trigger for GABAergic and AMPAergic scaling is distinct in mouse cortical cultures

We have shown the importance of alterations of spiking activity in triggering GABAergic scaling in mouse cortical cultures. Previously, we had shown that AMPAergic scaling was dependent on glutamatergic transmission rather than spiking, and did this in rat cortical cultures. This is a striking result as we had expected these homeostatic mechanisms to share a common trigger. To ensure that the triggers for AMPAergic and GABAergic scaling really were distinct in the same culture set and conditions used in the present study, we repeated our experiment blocking AMPAR activation for 24 hrs with 20µM CNQX, but now checked for AMPAergic scaling. We found the surprising result that following 24 hr CNQX treatment there was no change in AMPAergic mEPSC amplitudes (Figure 7), despite the fact that this was the same treatment that reduced spiking activity in our cultures and triggered GABAergic downscaling. The result confirms the observation that the triggers for AMPAergic and GABAergic scaling in the same cultures are distinct.

**Figure 7.**
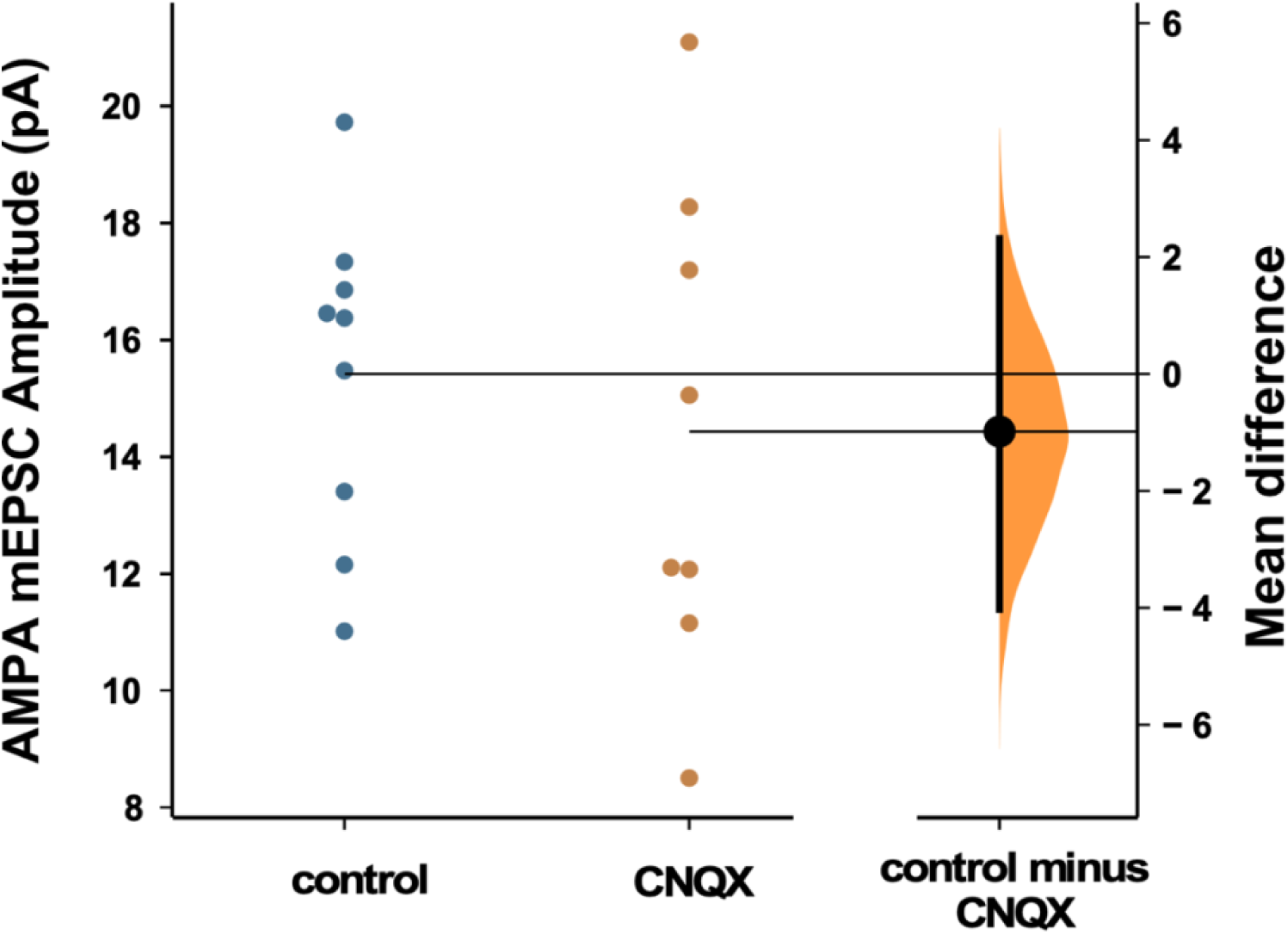
AMPAergic scaling was absent following 24 hours of 20µM CNQX. AMPA mEPSC amplitudes were no different than control following AMPAR blockade (p=0.57, control - n=9 from 4 cultures, CNQX - n=8 from 3 cultures). Recordings from single neurons (filled circles), where mean values (represented by horizontal bar) are plotted, as a bootstrap sampling distribution (mean difference is represented by a filled circles and the 95% CIs are depicted by vertical error bars). All mIPSC amplitudes recorded from cultures plated on MEAs.

## Discussion

In the original study describing AMPAergic synaptic scaling, the authors triggered this plasticity by blocking spiking activity with TTX or blocking AMPAergic neurotransmission with CNQX ^2^. Similar results have now been demonstrated in multiple tissues and labs ^26^. It was thought that AMPAergic scaling was a homeostatic mechanism, triggered by alterations in spiking and likely calcium transients associated with cellular spiking; once the cell drifted outside the setpoint for spiking a cell-wide signal was activated that changed the synaptic strengths of all AMPAergic inputs by a single multiplicative scaling factor to return the cell to the spiking set point ^3^. In this way, AMPAergic scaling could homeostatically regulate spiking levels, while also preserving the relative differences in synaptic strength that have been set up by Hebbian plasticity mechanisms. However, as described in the introduction, more recent studies suggest that AMPAergic synaptic scaling does not appear to meet these initial expectations.

Previous work suggests AMPAergic scaling following TTX or TTX+APV treatment was not multiplicative ^4, 5^, and we now show that it is not multiplicative following AMPAR blockade (CNQX treatment, Supplemental Figure 1). Further, several studies suggest that changes in mEPSC amplitude associated with AMPAergic scaling occur at the level of the synapse rather than globally throughout the cell. In fact, several studies have suggested that glutamate receptor activation due to action potential-independent spontaneous release could play a significant role in triggering AMPAergic scaling ^7, 11, 29^. It is certainly possible that there are compensatory changes in mEPSC amplitude that can be triggered by either altered neurotransmission or spiking. However, as we have shown previously, putting back significant spiking activity levels and their associated calcium transients in the presence of CNQX had no effect on AMPAergic scaling (no reduction in the existing scaling ^11^). Because AMPAergic scaling does not directly follow spiking activity levels, it does not appear to fulfill the expectations of a homeostat for spiking. Rather, AMPAergic scaling in many cases seems to act to homeostatically maintain the effectiveness of individual synapses.

GABAergic scaling appears to exhibit all the features initially predicted for AMPAergic synaptic scaling. First, GABAergic scaling is multiplicative, meaning the relative strengths of these synapses can be maintained (Figures 2, 4-6). Critically, GABAergic scaling can act as a firing rate homeostat for the following reasons. GABAergic downscaling was triggered by alterations in spike rate, rather than AMPAergic neurotransmission. We found that CNQX-triggered GABAergic downscaling was abolished when we optogenetically restored spiking activity levels (Figure 3-4), and that increasing spiking with bicuculline or CTZ both triggered GABAergic upscaling (Figures 5-6). While we cannot rule out a role of AMPAR activation in GABAergic upscaling, we did observe that CTZ-induced upscaling was converted to downscaling in the presence of TTX (Figure 5C-D). Further, the findings suggest that altering neurotransmission did not contribute to GABAergic scaling. Increasing AMPAergic transmission with CTZ in the presence of TTX had no impact on downscaling as it was no different than following TTX treatment alone (Figure 5D). Also, if GABAR transmission were a proxy for activity levels, then blocking GABA_A_ receptors would mimic activity blockade and should lead to a compensatory downscaling. However, bicuculline (reduced GABAR activity) increased spiking and triggered a GABAergic upscaling consistent with the idea that spiking was the critical feature (Figure 6). This result was consistent with previous work in hippocampal cultures where chronic bicuculline treatment triggered GABAergic upscaling, which was prevented if the cell was hyperpolarized ^18^. Finally, if scaling contributed to a homeostatic recovery of activity, then GABAergic scaling should have been expressed by 24 hours of CNQX (before bursts and before spike frequency in some cultures had fully recovered, Figure 1) and this was the case (Figure 2). Although AMPAergic scaling was initially thought to play the role of spiking homeostat, it appears more likely that GABAergic scaling is one of the homeostatic mechanisms that is playing this role.

The results of our current study on GABAergic scaling and our previous study on AMPAergic scaling ^11^ suggest these two forms of plasticity have distinct triggers and signaling pathways. Optogenetic restoration of activity in CNQX prevented GABAergic downscaling (Figure 3-4) but had no effect on AMPAergic scaling ^11^. Further, increasing glutamatergic receptor activation with CTX during activity blockade reduced TTX-induced AMPAergic scaling ^11^ but not GABAergic scaling (Figure 5D). We considered the possibility that some of our results could be due to differences in the cultures of this vs our previous study (mouse vs rat, 20 vs 40 µM CNQX, etc.). However, when we reduced spiking activity with 20µM CNQX and assessed AMPAergic scaling in mouse cortical cultures we did not trigger AMPAergic scaling at all, again consistent with the idea that the triggers are distinct for these two classes of plasticity. It is not clear to us why we were unable to trigger AMPAergic scaling in this study. It is possible that our cortical cultures (mouse, density) have less capacity for AMPAergic scaling. Alternatively, AMPAergic scaling may require higher concentrations of CNQX to partially influence NMDARs; this could occur through more complete blockade of AMPARs whose depolarization is important in removing the Mg block of the NMDAR or through direct block of the glycine binding site of the NMDAR ^30, 31^. Regardless, the reduction of spiking activity produced by 20µM CNQX was capable of triggering GABAergic scaling.

Previously, in embryonic motoneurons we found that both GABAergic and AMPAergic scaling was mediated by changes in GABAR activation from spontaneous release rather than changes in spiking activity ^12, 32^. However, this was at a developmental stage when GABA was depolarizing and could potentially activate calcium signaling pathways. On the other hand, spike rate homeostasis through the GABAergic system is consistent with many previous studies in which sensory input deprivation *in vivo* led to rapid compensatory disinhibition ^33, 34^. For instance, one day of visual deprivation (lid suture) reduced evoked spiking in fast spiking parvalbumin (PV) interneurons and this was thought to underlie the recovery of pyramidal cell responses to visual input at this point ^25^. One day of whisker deprivation between P17 and P20 produced a reduction of PV interneuron firing that was due to reduced intrinsic excitability in the GABAergic PV neuron ^21^. In addition, one day after enucleation, the excitatory to inhibitory synaptic input ratio in pyramidal cells was dramatically increased due to large reductions in GABAergic inputs to the cell ^24^. This disinhibition occurs rapidly ^23^ and can outlast changes in glutamatergic counterparts ^22, 24^. These results highlight the important role that inhibitory interneurons play in the homeostatic maintenance of spiking activity. Further, these cells have extensive connectivity with pyramidal cells, placing them in a strong position to influence network excitability ^35, 36^. In the current study, we show a critical feature of homeostatic regulation of spiking is through one aspect of inhibitory control, GABAergic synaptic scaling in excitatory neurons.

It is not clear what specific features of spiking triggers GABAergic scaling. GABAergic scaling may require the reduction of spiking in multiple cells in a network, rather than a single cell. Reduced spiking with sporadic expression of a potassium channel in one hippocampal cell in culture did not induce GABAergic scaling in that cell^17^. Such a result could be mediated by the release of some activity-dependent factor from a collection of neurons. BDNF is known to be released in an activity-dependent manner and has been shown to mediate GABAergic downward scaling following activity block previously in both hippocampal and cortical cultures ^16, 37^. On the other hand, another study increased spiking in hippocampal cultures and showed that homeostatic increases in mIPSC amplitudes could be dependent on the individual cell’s spiking activity ^18^. Future work will be necessary to determine the exact feature of spiking that may be more critical in triggering GABAergic scaling (e.g. bursting vs total spike frequency) and the downstream signaling pathway (e.g. somatic calcium transients).

While we have no direct support of a role for NMDARs, we cannot rule out the possibility that NMDAR activity could contribute to GABAergic scaling. Previous work has shown that NMDAR block can trigger GABAergic downscaling ^16^ and our activity manipulations would similarly alter NMDAR activation (CNQX would reduce and optogenetic restoration would restore some NMDAR activation). Whatever the specific features of spiking activity that trigger GABAergic scaling, our results strongly point to the idea that GABAergic scaling, could serve a critical role of a spiking homeostat, and highlights the fundamentally important homeostatic nature of GABAergic neurons.

Finally, it is important to take into consideration some of the benefits and limitations of this study. By recording activity levels of cultured neurons through multi-electrode arrays we were able to identify the actual influence of the drugs on population activity. This is a step beyond what many homeostatic studies, including our own, typically do, and it affords us the opportunity to interpret more intelligently the results of our perturbations. Regardless, there are limitations associated with these techniques. Cultured networks lack the actual circuitry of the *in vivo* cortex, and for several reasons are vulnerable to dramatic variability (based on plating density, ages, composition, etc.).

This variability can be seen in response to drug application throughout our results and it is important to keep in mind that the recorded spiking activity represents the population response from many different classes of excitatory and inhibitory neurons, although the majority are thought to represent excitatory principal neurons. Despite these caveats, the culture system has allowed us to manipulate spiking activity in important ways, which has provided us the insight that GABAergic scaling is one of the homeostatic mechanisms that fulfills the expectations of a spike rate homeostat.

## Materials and Methods

### Cell Culture

Brain cortices were obtained from C57BL/6J embryonic day 18 mice from BrainBits or harvested from late embryonic cortices. Neurons were obtained after cortical tissue was enzymatically dissociated with papain. Cell suspension was diluted to 2,500 live cells per ml and 35,000 cells were plated on glass coverslips or planar MEA coated with polylysine (Sigma, P-3143) and laminin. The cultures were maintained in Neurobasal medium supplemented with 2% B27 and 2mM GlutaMax. No antibiotics or antimycotics were used. Medium was changed completely after one day in vitro (1 DIV) and half of the volume was then changed every 7 days. Spiking activity was monitored starting ∼10 DIV to determine if a bursting phenotype was expressed and continuous recordings were made between 14-20 DIV. Cultures were discarded after 20 DIV. All protocols followed the National Research Council’s Guide on regulations for the Care and Use of Laboratory Animals and from the Animal Use and Care Committee from Emory University.

### Whole cell recordings

Pyramidal-like cells were targeted based on their large size. Whole-cell voltage clamp recordings of GABA mPSCs were obtained using an AxoPatch 200B amplifier, controlled by pClamp 10.1 software, low pass filtered at 5 KHz on-line and digitized at 20 KHz. Tight seals (>2 GΩ) were obtained using thin-walled boro-silicate glass microelectrodes pulled to obtain resistances between 7 and 10 MΩ. The intracellular patch solution contained the following (in mM): CsCl 120, NaCl 5, HEPES 10, MgSO_4_ 2, CaCl_2_ 0.1, EGTA 0.5, ATP 3 and GTP 1.5. The pH was adjusted to 7.4 with KOH.

Osmolarity of patch solution was between 280-300 mOsm. Artificial Cerebral-Spinal Fluid (ACSF) recording solution contained the following (in mM): NaCl 126, KCl 3, NaH_2_PO_4_ 1, CaCl_2_ 2, MgCl_2_ 1, HEPES 10 and D-glucose 25. The pH was adjusted to 7.4 with NaOH. GABAergic mPSCs were isolated by adding to ACSF (in µM): TTX 1, CNQX 20 and APV 50. Membrane potential was held at -70 mV and recordings were performed at room temperature. Series resistance during recordings varied from 15 to 20 MΩ and were not compensated. Recordings were terminated whenever significant increases in series resistance (> 20%) occurred. Analysis of GABA mPSCs was performed blind to condition with MiniAnalysis software (Synaptosoft) using a threshold of 5 pA for mPSC amplitude (50 mPSCs were taken from each cell and their amplitudes were averaged and each dot in the scatterplots represent the average of a single cell). Ratio plots of mIPSCs were constructed by taking a constant total number of mIPSCs from control and drug-treated cultures (e.g. 15 control cells with 40 mIPSCs from each cell and 20 CNQX-treated cells with 30 mIPSCs from each cell, 600 mIPSCs per condition). Then the amplitudes of mIPSCs from each condition were rank ordered from smallest to largest and plotted as a ratio of the drug-treated amplitude divided by the control amplitude, as we have described previously ^4, 26, 38^.

### MEA recordings

Extracellular spiking was recorded from cultures plated on planar 64 channel MEAs (Multichannel Systems) recorded between 14-20 DIV in Neurobasal media with B27 and GlutaMax, as described above. Cultured MEAs were covered with custom made MEA rings with gas permeable ethylene-propylene membranes (ALA Scientific Instruments). Synapse software (Tucker-Davis Technologies TDT) was used to monitor activity on a TDT electrophysiological platform consisting of the MEA MZ60 headstage, the PZ2 pre-amplifier and a RZ2 BioAmp Processor. Recordings were band-pass filtered between 200 and 3000Hz and acquired at 25KHz. MEA’s were placed in the MZ60 headstage, which was housed in a 5% C0_2_ incubator at 37°C. Drugs were added separately in a sterile hood and then returned to the MEA recording system. MEA spiking activity was analyzed offline with a custom-made Matlab program. The recordings acquired in Synapse software (TDT) were subsequently converted using the subroutine TDT2MAT (TDT) to Matlab files (Mathworks). The custom written Matlab program identified bursts of network spikes using an interspike interval-threshold detection algorithm ^39^. Spiking activity was labeled as a network burst when it met a user-defined minimum number of spikes (typically 10) occurring across a user-defined minimum number of channels (5-10) within a Time-Window (typically 0.1-0.3 seconds) selected based on the distribution of interspike intervals (typically between the first and 10th consecutive spike throughout the recording, Supplemental Figure 2). This program allowed us to remove silent channels and channels that exhibited high-noise levels. The identified network bursts were then visually inspected to ensure that these parameters accurately identified bursts. The program also computed network burst metrics including burst frequency, overall spike frequency and other characteristics.

### Optogenetic control of spiking

For photostimulation experiments neurons were plated on 64-channel planar MEAs and transfected with AAV9-hSynapsin–ChR2(H134R)-eYFP (ChR2) produced by the Emory University Viral Vector Core. All cultures used in ChR2 experiments, including controls, were transfected at 1 DIV. The genomic titer was 1.8×10^13^ vg/ml. Virus was diluted 1 to 10,000 in growth medium and this dilution was used for the first medium exchange at DIV 1. Finally, the media containing the virus was washed out after 24 hour incubation. A 3 hour pre-drug recording was obtained in the TDT program that determined the average MEA-wide firing rate before adding CNQX. This custom written program from TDT then delivered a TTL pulse (50-100ms) that drove a blue light photodiode (465 nm, with a range from 0 to 29.4 mwatts/mm^2^, driven by a voltage command of 0-4V) from a custom-made control box that allowed for scaled illumination. The photodiode sat directly below the MEA for activation of the ChR2. This triggered a barrage of spikes resulting in a burst that looked very similar to a naturally occurring burst not in the presence of CNQX. The program measured the MEA-wide spike rate every 10 seconds and if the rate fell below the set value established from the pre-drug average, an optical stimulation (50-100ms) was delivered triggering a burst which then increased the average firing rate, typically above the set point.

### Statistics

Estimation statistics have been used throughout the manuscript. 5000 bootstrap samples were taken; the confidence interval is bias-corrected and accelerated. The *P* value(s) reported are the likelihood(s) of observing the effect size(s), if the null hypothesis of zero difference is true. For each permutation *P* value, 5000 reshuffles of the control and test labels were performed (Moving beyond P values: data analysis with estimation graphics ^40^).

## Acknowledgments

We would like to thank Bill Goolsby who custom built our optogenetic stimulator, and Tucker Davis Technologies for helping us write the Synapse Program that ran the MEA recording/optogenetic stimulation software. We would also like to thank Dr. Gary Bassell for providing us with some of the mice used in culture experiments.

---

**Supplemental Figure 1.**
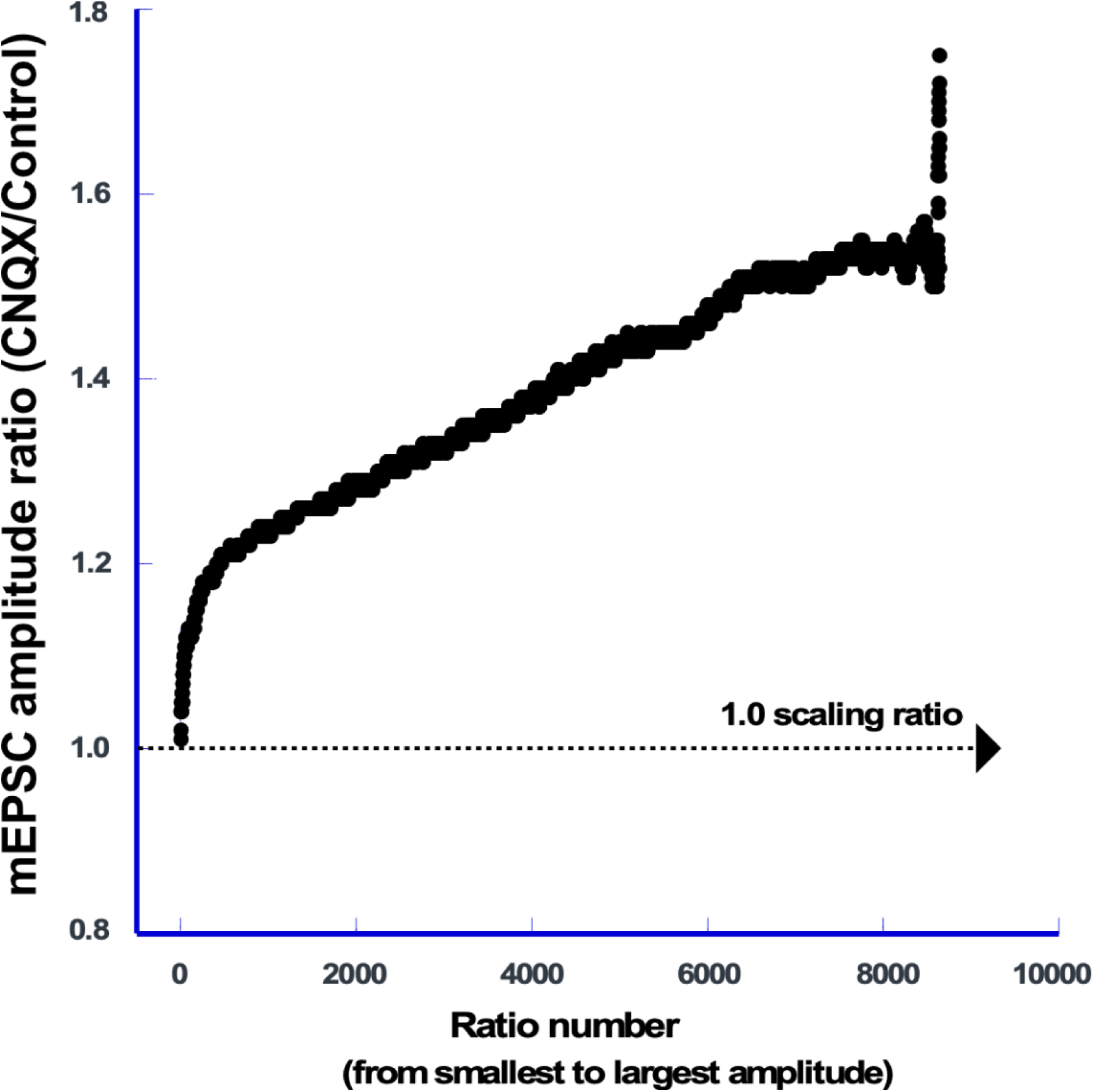
AMPAR block triggered non-uniform AMPAergic scaling. Scaling ratio plot shows the ratio of rank ordered mEPSC amplitudes from CNQX-treated cultures (n=95 cells, 91mEPSCs/cell) divided by those from untreated cultures (n=91 cells, 95 mEPSCs/cell). The X axis represents the rank ordered number of mEPSCs (from smallest to largest).

**Supplemental Figure 2.**
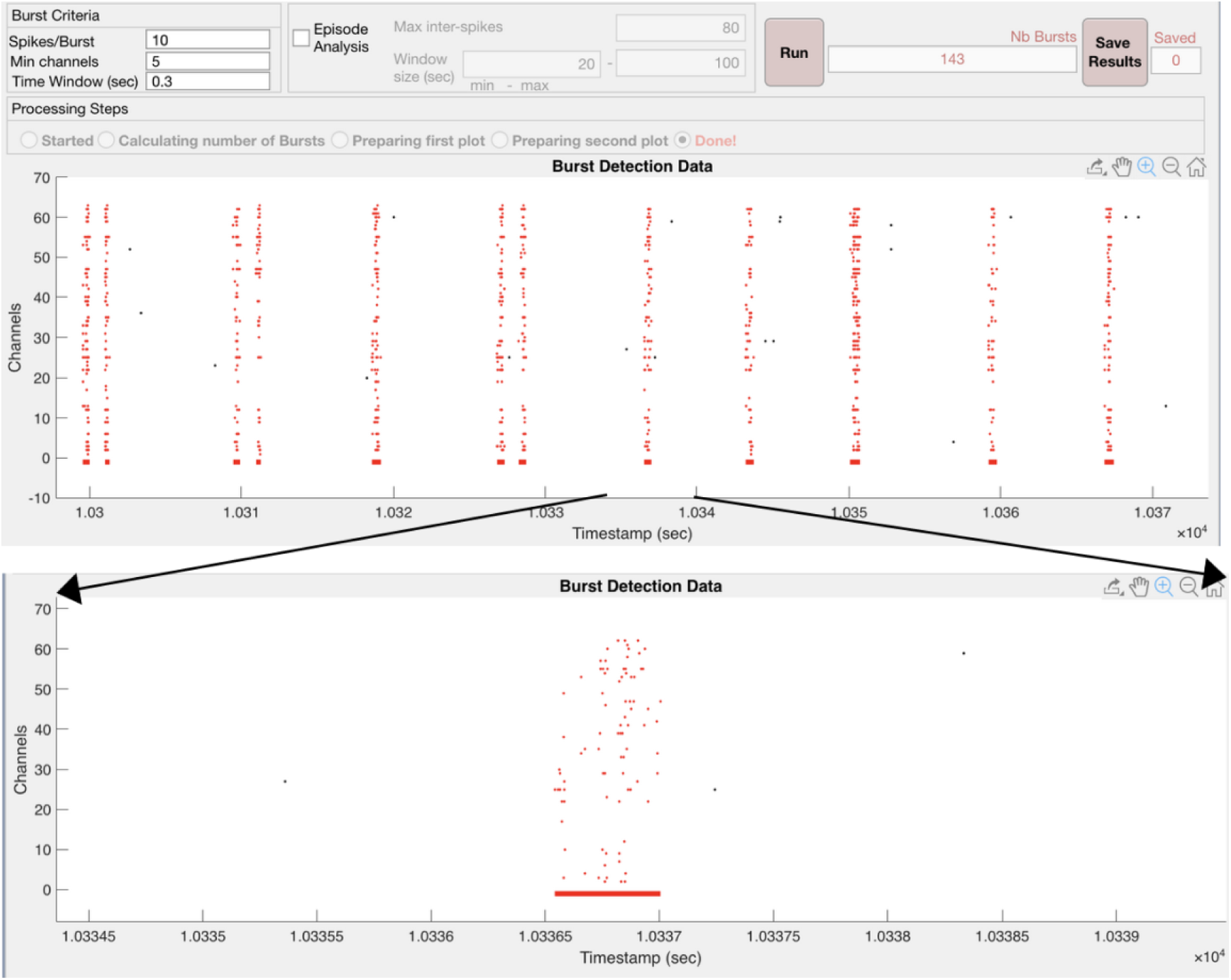
Custom written Matlab program identifies bursts in cortical cultures plated on MEA’s by choosing the minimum number of spikes per burst (Spikes/Burst) across a minimum number of channels contributing to a burst (Min channels) within a maximum Time Window. Upper image shows the identification of bursts in red across 64 channels as a raster plot where each dot represents one spike detected on the MEA. The program then examines various parameters which were then exported to an excel spreadsheet for analysis. Burst identity and duration are shown as a red line positioned below the raster plot. A single burst is expanded and plotted below the upper image.

**Supplemental Figure 3.**
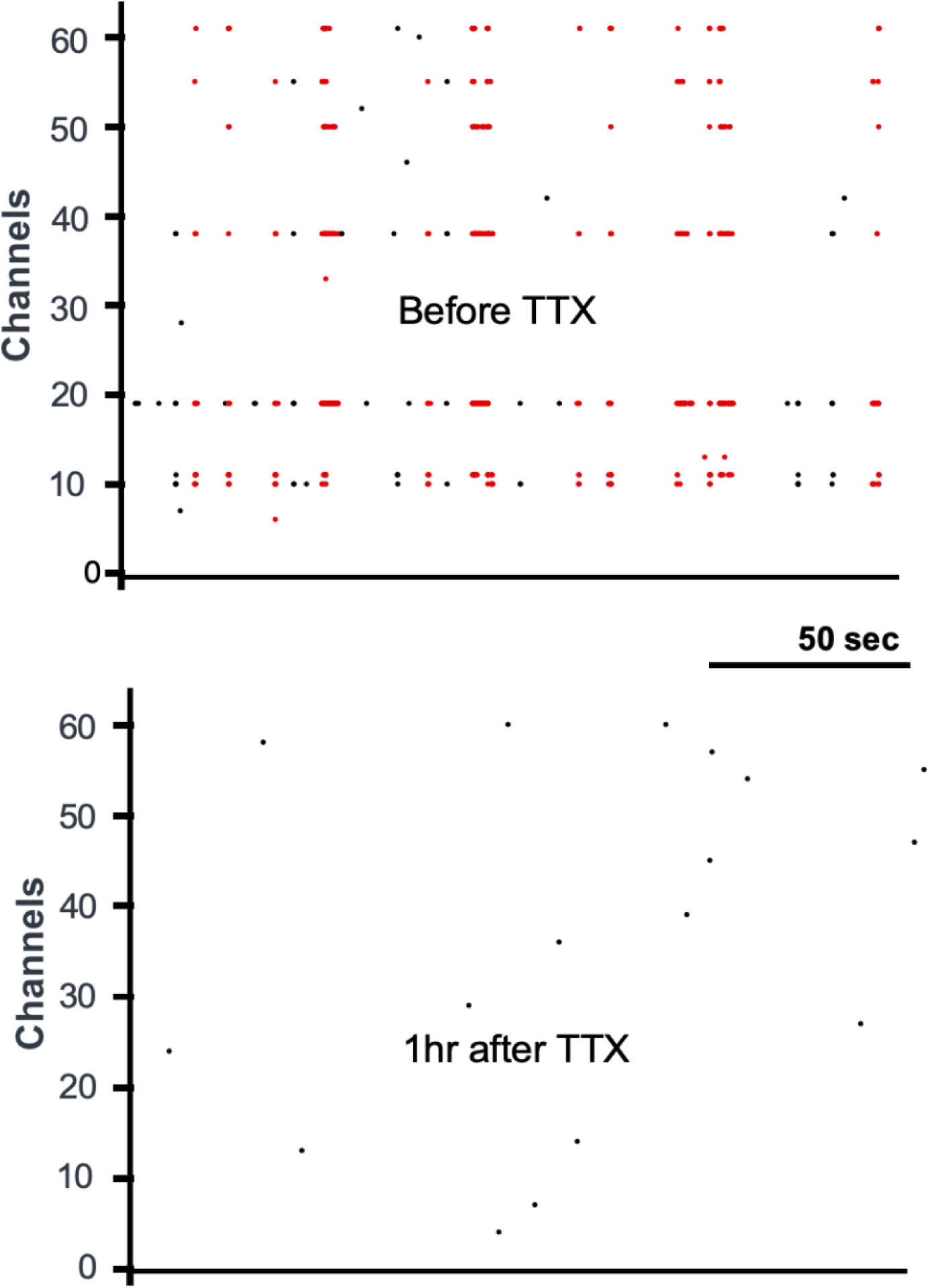
Raster plot of cortical culture plated on MEA demonstrating network bursting (red dots, upper plot). Bursts were then abolished after addition of TTX (1µM) to the culture; a small number of spike detections remain, however these are likely to be noise that crosses the detection threshold.

**Supplemental Figure 4.**
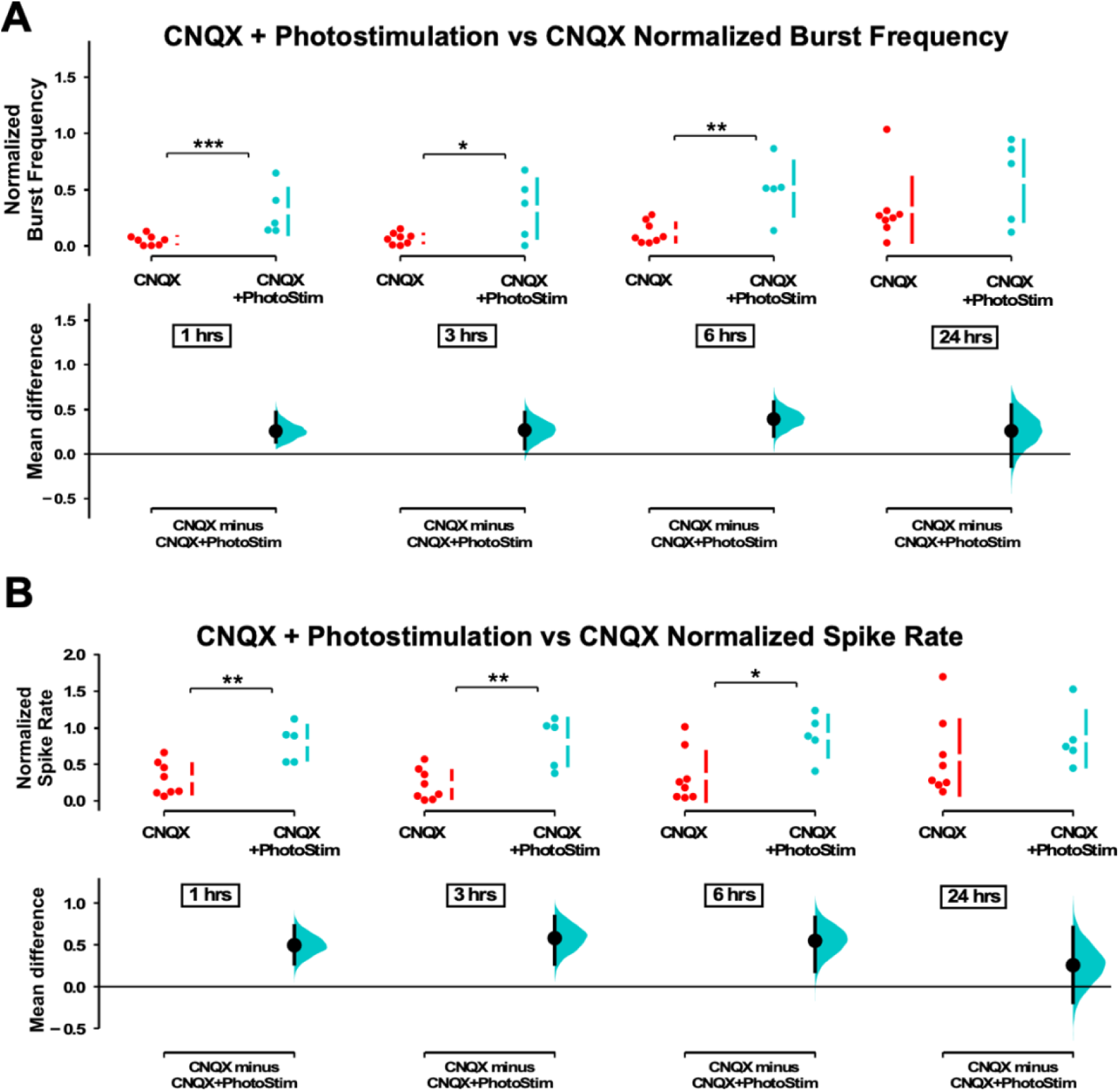
MEA recording’s show optostim + CNQX increases burst frequency and spike frequency compared to CNQX alone. **A**) Average burst rate is compared for CNQX-treated cultures with optogenetic stimulation (n=5) and CNQX only unstimulated cultures (n=8) at 1hr, 3hrs, 6hrs, and 24hrs (p=0.209) after addition of CNQX. **B**) Average overall spike frequency is compared for CNQX-treated cultures with optogenetic stimulation and CNQX only unstimulated cultures at 1hr, 3hrs, 6hrs, and 24hrs (p=0.389) after addition of CNQX. The mean differences at different time points are compared to control and displayed in Cumming estimation plots. Significant differences denoted by * p ≤ 0.05, ** p ≤ 0.01, *** p ≤ 0.001. Recordings from single cultures (filled circles), where mean values (represented by the gap in the vertical bar) and SD (vertical bars) are plotted on the upper panels. Mean differences between control and treated groups are plotted on the bottom panel, as a bootstrap sampling distribution (mean difference is represented by a filled circles and the 95% CIs are depicted by vertical error bars).

**Supplemental Figure 5.**
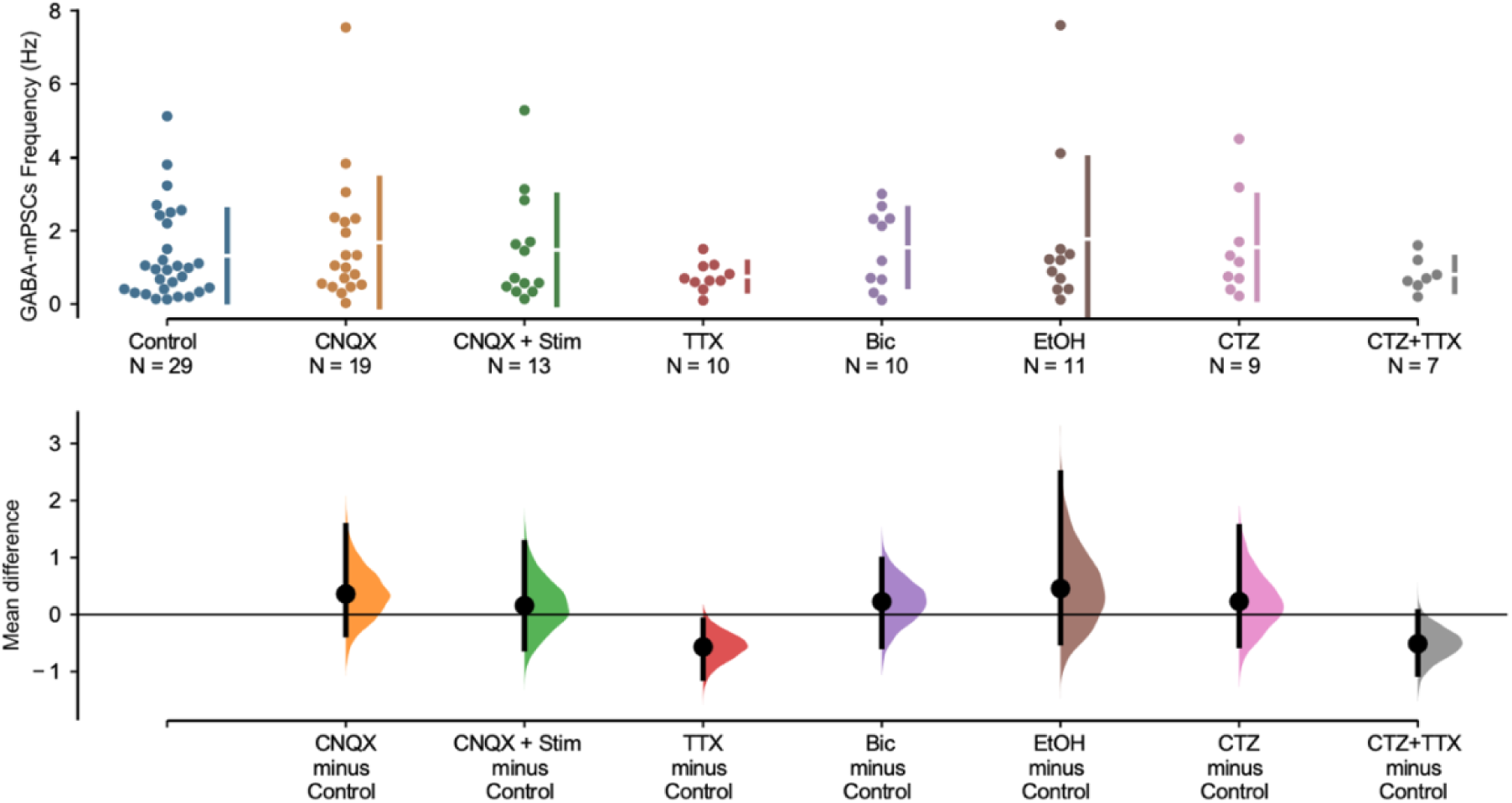
Frequency of mIPSCs were no different across conditions. Scatterplots of mIPSC frequency show tremendous variability but do not exhibit significant differences through different drug treatments. The mean differences are compared to control and displayed in Cumming estimation plots. GABAergic mPSC frequencies from single neurons (filled circles), where mean values (represented by the gap in the vertical bar) and SD (vertical bars) are plotted on the upper panels. Mean differences between control and treated groups are plotted on the bottom panel, as a bootstrap sampling distribution (mean difference is represented by a filled circles and the 95% CIs are depicted by vertical error bars).

## Notes

**Competing Interest Statement:** We have no competing interests.

### Competing Interest Statement

The authors have declared no competing interest.

### Summary of Updates

This version of the manuscript has been revised to respond to the reviewers' comments at eLife. This includes changes throughout the text of the mauscript and figures, including an additional experiment and associated figure.

